# The role of *miR-9a* in modulating sensory neuron morphology and mating behavior in *Drosophila melanogaster*

**DOI:** 10.1101/2025.09.09.675227

**Authors:** Tianmu Zhang, Hongyu Miao, Xiaoli Zhang, Joshua Bagley, Yongwen Huang, Woo Jae Kim

**Author notes:** These authors contributed equally (Tianmu Zhang, Hongyu Miao). AUTHOR CONTRIBUTIONS TZ and HM designed and performed revision experiments and revised manuscript. JB designed and performed experiments. XZ and YH supported additional revision experiments. WJK designed and performed experiments, wrote, and revised manuscript, analyzed data, and performed image analysis.

## Abstract

Following mating, female *Drosophila melanogaster* display profound behavioral changes, including sensory sensitivity and rejection of courting males. The molecular mechanisms governing this plasticity remain incompletely understood. Here, we identify the conserved microRNA, miR-9a, as a critical regulator of this process. We show that miR-9a mutant females exhibit a premature rejection phenotype, mimicking mated-female behavior, which is correlated with an aberrant overgrowth of adult body wall sensory neurons. We demonstrate that this neuronal phenotype is governed by a dual regulatory system. First, in a non-cell autonomous mechanism, miR-9a expression in the epidermis is required to constrain sensory neuron dendrite growth, indicating that an epithelial-derived signal patterns the underlying neuron. Second, within the neuron itself, miR-9a interacts genetically with the transcription factor *senseless* (*sens*) and the novel RNA-binding protein *bruno2 (bru2)*. Reducing the dosage of either *sens* or *bru2* rescues both the neuronal and behavioral defects of miR-9a mutants. Our findings reveal an integrated, inter-tissue signaling axis where epithelial miR-9a orchestrates a non-cell autonomous cue that modulates a cell-intrinsic network to ensure the precise development of sensory neurons, thereby calibrating behavioral responses critical for reproductive success.

## INTRODUCTION

After mating, female *Drosophila melanogaster* undergo a marked transformation in behavioral response, exhibiting an enhanced sensitivity to a broad array of sensory inputs. This altered sensory perception is a component of a complex suite of post-mating changes that optimize the female’s reproductive strategy (Yang et al. 2009a; Zhu et al. 2014; Hussain et al. 2016; Bath et al. 2017). Enhanced sensory reactivity represents a pivotal component of the post-mating behavioral syndrome in female *Drosophila melanogaster*. This adaptive sensitization is hypothesized to confer selective advantages by facilitating the evasion of predators and/or by aiding in the detection of optimal oviposition sites, thereby enhancing the female’s reproductive success (Hollis et al. 2019).

MicroRNAs (miRNAs) are a class of endogenous small non-coding RNAs that predominantly function in the post-transcriptional regulation of gene expression. These molecules exert their regulatory effects by annealing to the complementary sequences within the 3’ untranslated region (3’ UTR) of target messenger RNAs (mRNAs), thereby facilitating either the degradation of the mRNA transcript or the suppression of protein synthesis, depending on the degree of complementarity and other context-dependent factors (Bartel 2009a; Brodersen and Voinnet 2009). miRNAs are implicated in diverse brain functions including development, cognition, and synaptic plasticity (Smalheiser and Lugli 2009a; Cohen et al. 2011a; Aksoy-Aksel et al. 2014a; Ye et al. 2016; Mohammadi et al. 2022a).

The *miR-9a* and its mammalian ortholog, *miR-9*, are pivotal regulators of post-transcriptional gene expression, with diverse roles in development, cellular differentiation, and disease progression (Li et al. 2006; Biryukova et al. 2009; Cassidy et al. 2013a; Li et al. 2013; Yatsenko and Shcherbata 2014; Cassidy et al. 2015; Suh et al. 2015; Daniel et al. 2017; Katti et al. 2017; Gallicchio et al. 2020; Subramanian et al. 2021). Although *miR-9a* is multifunctional, *miR-9a* was initially recognized for its essential function in neural development, particularly in the precise specification of sensory organ precursors (SOPs) (Li et al. 2006; Parrish et al. 2006). Notably, *miR-9a* exhibits a strong genetic interaction with the *senseless* (*sens*) gene in controlling SOPs formation (Li et al. 2006). The *miR-9a* predominantly influences the development of multidendritic sensory neurons and is crucial for the correct morphogenesis of their dendritic arbors (Parrish et al. 2006; Wang et al. 2016a).

In this study, we investigated the role of *miR-9a* in regulating reproductive behaviors and neuronal development in *Drosophila melanogaster*. We found that virgin females carrying *miR-9a* mutations display reduced receptivity, increased rejection of courting males, and abnormal mating termination behavior, while males show distinct defects in courtship displays. These behavioral changes correlate with aberrant overgrowth of body wall sensory neurons, suggesting that *miR-9a* influences pre-mating behaviors through regulation of neuronal development. Furthermore, genetic interaction experiments with *sens* and the newly identified interactor *bru2* revealed that partial loss of these genes rescues the *miR-9a* mutant phenotype. Together, our findings demonstrate that *miR-9a* plays a crucial role in sensory neuron specification and pre-mating reproductive behaviors in *Drosophila*.

## RESULTS

### Female *Drosophila melanogaster* with mutations in *miR-9a* exhibit rejection of courting males

It is established that internal sensory neurons expressing the *pickpocket* (*ppk*) gene mediate the post-mating behavioral switch through the binding of sex peptide to its receptor (Häsemeyer et al. 2009a; Yang et al. 2009a). Given that *miR-9a* is implicated in the development of sensory organ precursors (SOPs) (Li et al. 2006; Wang et al. 2016a), including *ppk*-positive neurons, indicating that *miR-9a* mutations are associated with altered receptivity and may involve changes in *ppk*-expressing neurons.

We used two previously characterized loss-of-function alleles of *miR-9a*, *miR-9a^J22^* and *miR-9a^E58^*. Both alleles carry deletions that remove the precursor hairpin sequence of *miR-9a*, resulting in a complete loss of mature *miR-9a* expression (Li et al. 2006; Biryukova et al. 2009). These alleles have been widely used to study *miR-9a*’s role in neural development and sensory organ specification. Both *miR-9a* mutant strains, *miR-9a^J22^*and *miR-9a^E39^*, displayed a significant decrease in receptivity among virgin females across both 20-minute and 120-minute observation periods (Fig. 1A-B, Fig. S1A-B and Fig. S1F-G). Temporal analysis of receptivity scores definitively revealed that *miR-9a* mutant females exhibit a substantial delay in the onset of sexual receptivity (Fig. 1C and Fig. S1C). Relative to wild-type control females, *miR-9a* mutant females exhibited a significantly higher rejection rate within a 5-minute observation window (Fig. 1D). Moreover, *miR-9a* mutant females that initially accepted a male often subsequently terminated the mating attempt (Fig. 1E). This behavior, wherein a female accepts a male and then rapidly rejects accepted male within seconds, is a phenotype that is infrequently observed in wild-type females (Movies 1-4).

**Figure 1.**
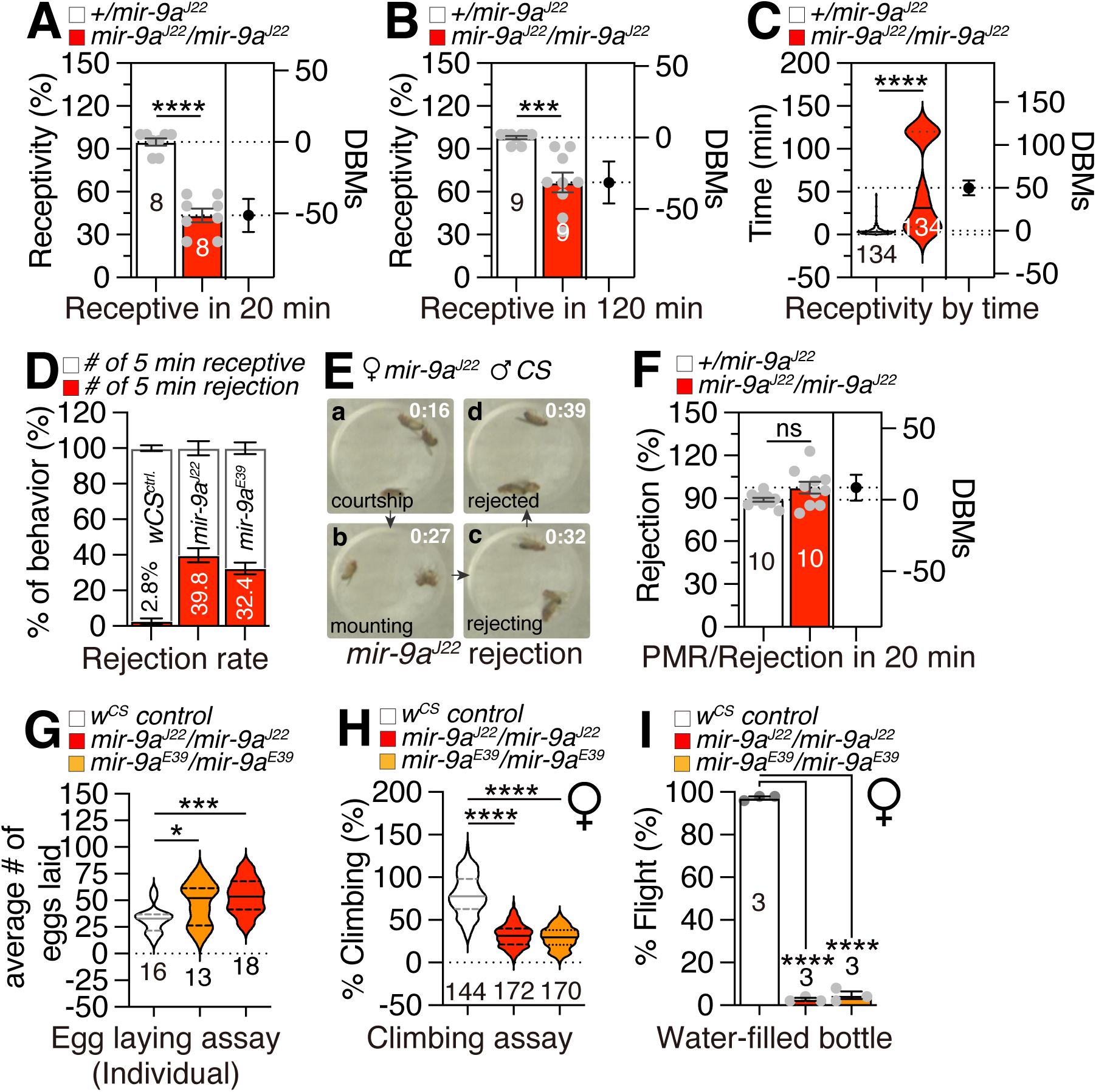
*miR-9a* mutations lead to a specific increase in rejection behavior among virgin females. (A-B) Receptivity of virgin females, with the score for receptivity behavior as well as genotype of experimental animals given above the graph. In each assay, one female of the indicated genotype was confronted with two naive males in a small chamber. Females were scored as receptive when they mated within 20 minutes (A) and 120 minutes (B). Numbers in columns are the numbers of females tested. DBMs represent difference between means. Error bars show the 95% confidence interval for the mean difference, calculated using estimation statistics. The mean value and standard error are labeled within the dot plot (black lines). Asterisks represent significant differences, as revealed by the Student’s t test (* p<0.05, ** p<0.01, *** p<0.001). The same notations for statistical significance are used in other figures. (C) Temporal dynamics of virgin female receptivity over a 120-minute observation period. Each curve represents an independent time-course experiment performed separately from the receptivity assays shown in Figures 1A and 1B, which explains the difference in sample sizes. Females were scored as receptive when copulation occurred, and cumulative receptivity was plotted across the indicated time intervals. Numbers in parentheses denote the number of females tested per genotype. (D) The Receptivity assays of *wCS* females, *miR-9a^J22^/miR-9a^J22^* homozygous mutant and *miR-9a^E39^/miR-9a^E39^* homozygous mutant in 5 minutes. (E) Females harboring a *miR-9a^J22^* mutation initially accepted male courtship but subsequently terminated the mating attempt. (F) Rejection of mated females, with the score for rejection behavior as well as genotype of experimental animals given above the graph. PMR means post-mating reaction. For detailed description of the calculation of rejection rate, see **Receptivity Assay** in **Method section** (G) Egg laying. For egg laying, 10 females of the appropriate genotype were aged in vials for 4–5 days. Then three or five females were transferred to a vial with grape media and allowed to lay eggs for 24 hr at 25°C. The number of eggs was divided by the number of flies in the vial to give a measure of egg laying. (H) Climbing assays of females. 40–50 flies were placed in an empty vial and were tapped to the bottom of the tube. After tapping of flies, we recorded 10 s of video clip. This experiment was done five times at 5-min intervals. With recorded video files, we captured the position of flies 10 s after tapping the vial. (I) Flight assays of females. For each assay, fifty flies were gently introduced into a water-filled jar. The jar was then tapped to stimulate the flies, and the number of flies that escaped from the water was counted. The percentage (%) of flies that escaped was calculated to determine the effectiveness of the flies’ flight response.

Post-mating rejection of male courtship is a conserved element of female post-mating responses (PMR) in *Drosophila melanogaster* (Chapman et al. 2003; Kubli 2003; Yapici et al. 2008). While *miR-9a* mutants did not show a detectable difference in post-mating rejection compared with controls, the rejection rate of mated *miR-9a* mutants was indistinguishable from that of control mated females (Fig. 1F and Fig. S1D), indicating that the PMR in *miR-9a* mutants is intact. Notably, mated *miR-9a* mutants deposited a greater number of eggs than controls (Fig. 1G and Fig. S1E), suggesting that *miR-9a* specifically modulates egg-laying behavior rather than rejection behavior within the PMR. Furthermore, *miR-9a* mutants exhibited a marked reduction in climbing (Fig. 1H) and flight (Fig. 1I; Movies 5-6) abilities, indicative of impaired muscle contraction, consistent with previous reports (Katti et al. 2017). Collectively, these findings indicate that *miR-9a* mutations lead to a specific increase in rejection behavior among virgin females, while the rejection responses of mated females remain unaffected.

### Male fruit flies with mutations in *miR-9a* display aberrant courtship behaviors, and the larvae of these mutants exhibit locomotor defects

Although our main focus is on virgin female receptivity, we also noted that *miR-9a* mutant males exhibit altered courtship performance. Specifically, mutant males showed a reduced courtship index (Fig. 2A). Notably, a substantial proportion of *miR-9a* mutant males displayed a unique courtship behavior characterized by bilateral wing vibration (Fig. 2B), in contrast to the typical unilateral wing vibration observed in controls (Fig. 2C). Approximately two-thirds of *miR-9a* mutant males exhibited this double wing vibration phenotype (Fig. 2D and Movies 7-8), indicating that *miR-9a* mutation leads to specific courtship defects in male flies. Similar to the locomotor deficits observed in mutant females, *miR-9a* mutant males also exhibited reduced climbing and flight abilities (Fig. S2A-B), suggesting a generalized impact of *miR-9a* mutation on locomotor behavior. These secondary results suggest that *miR-9a* influences reproductive behaviors in both sexes, though with distinct outcomes.

**Figure 2.**
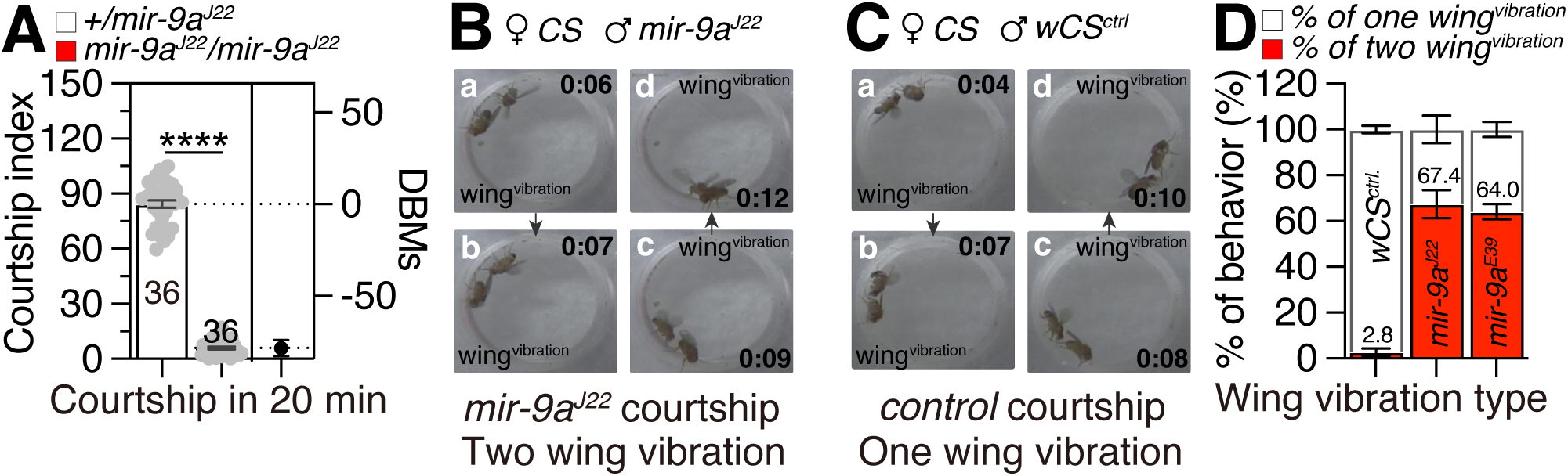
Males with *miR-9a* mutation display abberent courtship behaviors. (A) Courtship assays of males. Once courtship began, courtship index was calculated as the fraction of time a male spent in any courtship-related activity during a 10 min period or until mating occurred. Mating initiation is the time after male flies successfully mounted on female. (B) *miR-9a^J22^* males displayed a unique courtship behavior characterized by bilateral wing vibration. (C) *CS* males displayed a typical courtship behavior characterized by unilateral wing vibration. (D) Wing vibration types of males. White box represents the percentage of unilateral wing vibration and red box represents the percentage of bilateral wing vibration.

In line with these adult locomotion data, third instar larvae of *miR-9a* mutants showed impaired locomotion (Fig. S2C-E) and exhibited an increased frequency of turning behaviors compared to controls (Fig. S2F and Movies 9-10), indicating that the *miR-9a* mutation elicits a unique behavioral profile, distinct from a generalized impairment of health. These findings collectively suggest that *miR-9a* mutation induces a range of sensory-motor-related phenotypes, from larvae to adults, across both sexes.

### In *miR-9a* mutant flies, the adult body wall neurons undergo aberrant overgrowth

The conservation of miR-9 across evolutionary scales, from flies to humans, at the nucleotide level underscores its functional significance, despite variations in expression patterns (Leucht et al. 2008; Biryukova et al. 2009; Coolen et al. 2013) . Studies across various model organisms have revealed that miR-9 can influence neurogenesis through its regulatory role in both neural and non-neural cell lineages (Yuva-Aydemir et al. 2011a). In *Drosophila*, *miR-9a* is essential for the accurate specification of neural progenitor cells in non-neural lineages. For instance, the loss of *miR-9a* activity leads to the formation of ectopic sensory neurons in embryos, larvae, and adults (Li et al. 2006).

To assess whether the female mating rejection phenotype is associated with aberrant sensory neuron development, we analyzed the female abdominal body wall neurons, which derive from the larval sensory neurons that are regulated by *miR-9a* (Li et al. 2006). Consistent with previous findings, *ppk*-positive larval body wall sensory neurons in *miR-9a* mutants exhibit ectopic sensory neurons with exceptionally branched dendrites (Fig. S3A-B). Remarkably, adult body wall neurons also display ectopic sensory neurons with overgrown dendrites in both the ventral and dorsal abdomen (Fig. 3A-B and Fig. S3C-D). Quantification of neurite morphology revealed a significant increase in the number of branches and junctions in *miR-9a* mutant body wall neurons (Fig. 3C-D and Fig. S3E-F), although the average length of the branches is comparable between the mutant and control groups (Fig. 3E and Fig. S3G). These data show that *miR-9a* mutants display aberrant adult body wall neuronal growth, consistent with its known roles in neuronal specification.

**Figure 3.**
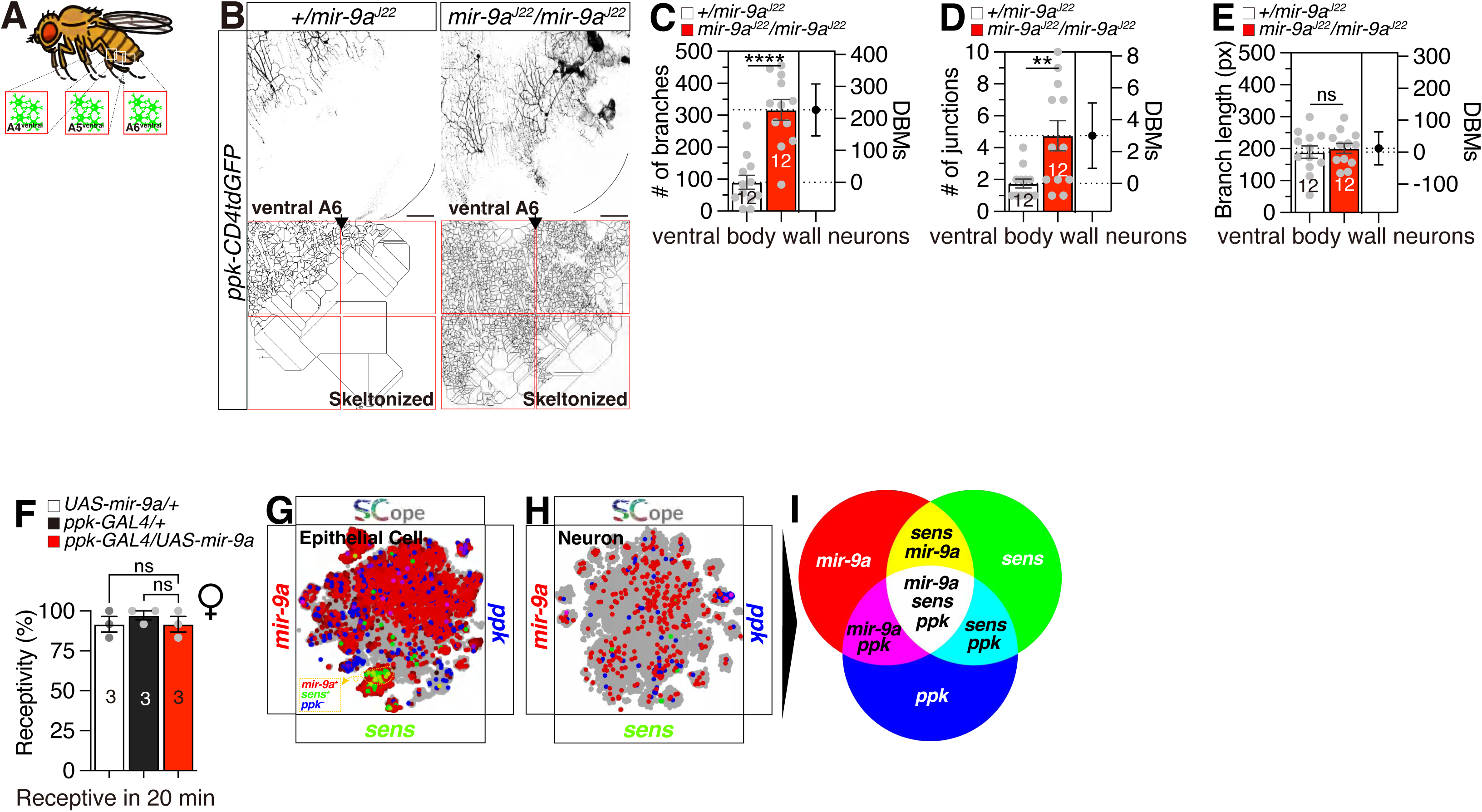
The adult ventral body wall neurons undergo aberrant overgrowth in *miR-9a* mutant flies. (A) Location of *Drosophila* adult ventral neurons. (B) Ventral A6 neurons expressing *ppk-GAL4* together with *UAS-mCD4GFP* in *miR-9a^J22^/ +* and *miR-9a^J22^/miR-9a^J22^* in adult female. The bottom pictures are skeletonized from the top pictures. Scalebar represent 100 μm. (C-E) Quantification of neurite morphology for *miR-9a* mutant body wall neurons in branches (C), junctions (D) and branch length (E). (F) Receptivity of virgin females in 20 minutes, with the score for receptivity behavior as well as genotype of experimental animals given above the graph. (G-H) Single-cell RNA sequencing (SCOPE scRNA-seq) datasets reveal tissues colored by expression of *miR-9a* (red), *ppk* (blue) with *sens* (green). (I) Schcematic diagram for color code presented in (G-H).

### *miR-9a* does not act cell-autonomously within *ppk^+^* neurons

To test for a cell-autonomous requirement of *miR-9a* in *ppk^+^* neurons, we first overexpressed it in these cells using the *ppk-GAL4* driver. This manipulation had no discernible effect on female receptivity compared to controls, arguing against a simple model where *miR-9a* levels within ppk+ neurons directly dictate mating behavior (Fig. 3F).

To further probe the spatial relationship between *miR-9a* and its interactors, we analyzed publicly available single-cell RNA sequencing data (Fly Atlas SCope) (Li et al. 2022). This analysis revealed no significant co-expression between *miR-9a* and the *ppk* marker in relevant cell clusters. We then examined the expression of *senseless* (*sens*), a known co-factor involved in sensory neuron’s function. As previously reported, *miR-9a* and *sens* were found to co-express highly, but this overlap was restricted to epithelial cells (Fig. 3G). Critically, we could not find any clear co-expression between *miR-9a* and *ppk*, or between *miR-9a* and *sens*, within any neuronal populations identified in the dataset (Fig. 3H-I).

Taken together, these genetic and transcriptomic data strongly suggest that *miR-9a* does not function directly within *ppk^+^* neurons. The lack of a behavioral phenotype from neuronal-specific overexpression, combined with the spatial separation of its expression from both *ppk* and neuronal *sens*, supports a non-cell-autonomous mode of action. This model, which aligns with previous work (Li et al. 2006; Biryukova et al. 2009; Cassidy et al. 2013b; Wang et al. 2016b; Gallicchio et al. 2020), suggests *miR-9a* likely acts in other tissues, such as the epithelium, to indirectly influence the function of *ppk^+^* neurons and modulate female mating behavior.

### An epithelial *miR-9a/sens* interaction non-cell autonomously regulates sensory neuron morphology and behavior

It has been reported *miR-9a* mutants exhibit sensory bristle defects on the notum, and *miR-9a* interacts genetically with the *sens* gene (Li et al. 2006). To determine whether the effects of *miR-9a* mutation on adult body wall neurons involve interaction with the *sens* gene, we conducted genetic interaction experiments. In a *miR-9a^J22^* homozygous mutant background, the removal of one copy of *sens* (*sens^E58^/+*) resulted in a significant rescue of the receptivity defects, matching those of control females (Fig. 4A and Fig. S4A). This genetic rescue of the receptivity phenotype is correlated with the normalization of the adult body wall neuronal phenotype, which displays normal sensory neurons with typical neurite growth (Fig. 4B-E). We did not observe differences in the overall number of *ppk^+^* sensory neurons in the abdominal body wall (A4–A6) between controls and *miR-9a* mutants, indicating that the overgrowth phenotype reflects increased branching within existing neurons rather than changes in neuron number (Fig. S4B).

**Figure 4.**
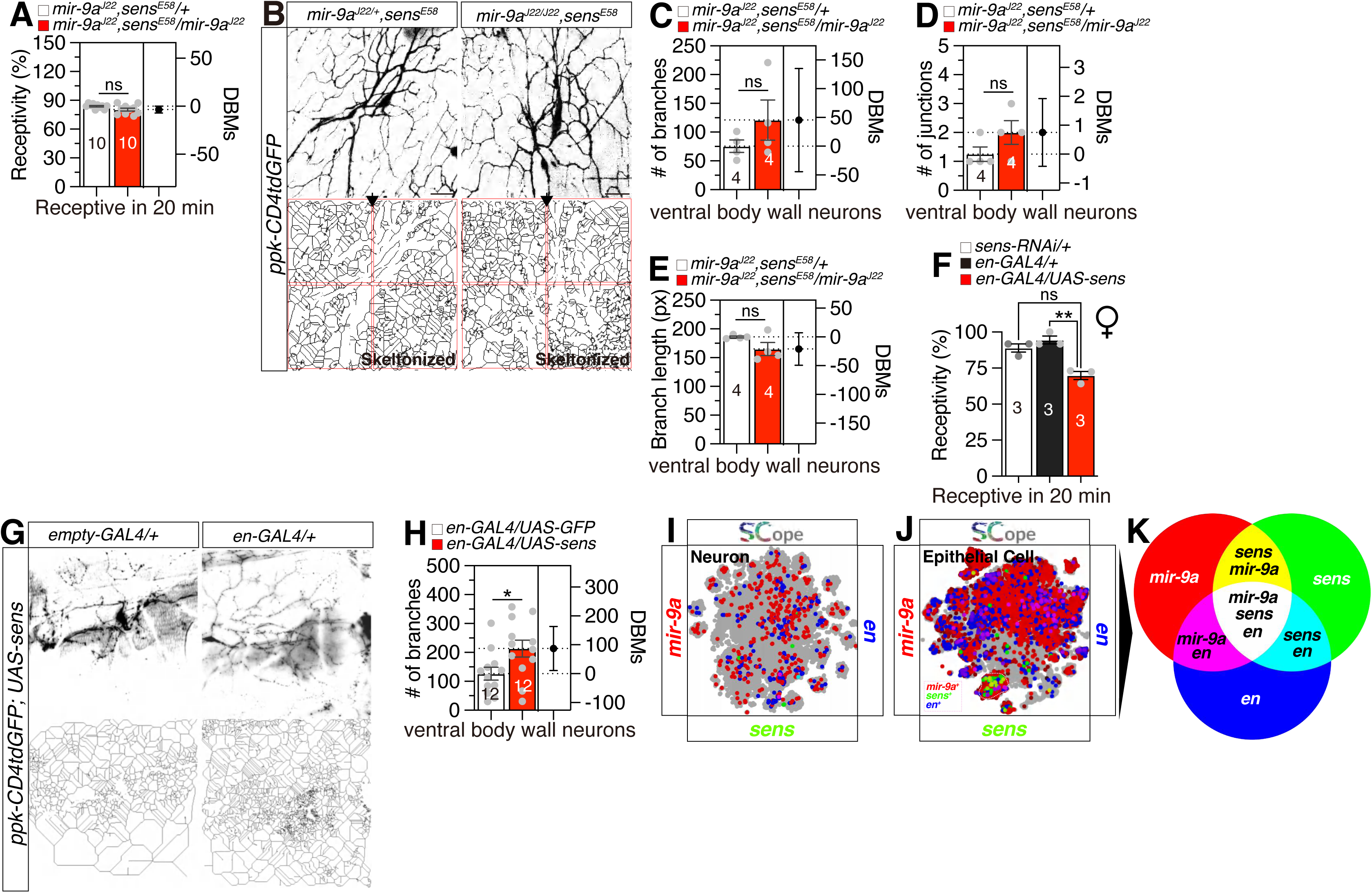
*sens* and *bru2* mutations rescue the phenotypic defects of *miR-9a* mutant flies. (A) Receptivity of virgin females, with the score for receptivity behavior as well as genotype of experimental animals given above the graph. (B) Ventral neurons expressing *ppk-GAL4* together with *UAS-mCD4GFP* in *miR-9a^J22^/ +, sens^E58^* and *miR-9a^J22^/miR-9a^J22^, E58* female. The bottom pictures are skeletonized from the top pictures. Scalebar represent 10 μm. (C-E) Quantification of neurite morphology for venral body wall neurons in branches (C), junctions (D) and branch length (E). Genotype of experimental animals given above the graph. (F) Receptivity of virgin females in 20 minutes, with the score for receptivity behavior as well as genotype of experimental animals given above the graph. (G) Overexpression of *sens* in body wall sensory neurons via *en-GAL4* with *ppk-CD4tdGFP*. (H) Quantification of neurite morphology for body wall sensory neurons in branches. Genotype of experimental animals given above the graph. (I-K) Single-cell RNA sequencing (SCOPE scRNA-seq) datasets reveal tissues colored by expression of *miR-9a* (red), *en* (blue) with sens (green). (K) Schcematic diagram for color code presented in (I-K).

To elucidate the non-cell autonomous mechanism by which *miR-9a* influences sensory neurons, we tested whether its regulatory effect is mediated through *sens* expression in the adjacent epithelium. We hypothesized that manipulating *sens* levels exclusively in epithelial cells would be sufficient to replicate the neuronal and behavioral phenotypes observed in *miR-9a* mutants. To test this, we employed the *en-GAL4* driver to specifically overexpress a *UAS-sens* transgene in the epithelium (Brower 1986; Blagburn 2008).

This epithelium-specific overexpression of *sens* successfully phenocopied the *miR-9a* mutant. We observed a significant increase in the dendritic branching of *ppk*-positive body wall sensory neurons (Fig. 4G-H), mirroring the morphological defects of the *miR-9a* loss-of-function. Furthermore, this genetic manipulation produced a corresponding behavioral deficit, manifesting as significantly reduced sexual receptivity in female flies (Fig. 4F).

To confirm the spatial segregation of these components, we analyzed fly SCope single-cell RNA sequencing data. The analysis revealed that miR-9a, *sens*, and *en* transcripts are highly co-expressed within a distinct cluster of epithelial cells, separate from the neuronal clusters expressing *ppk* (Fig. 4I-K, cf. Fig. 3G-I). This lack of overlap provides compelling evidence that the *en-GAL4*-mediated overexpression of *sens* in the epithelium affects sensory neuron morphology through a non-cell autonomous pathway.

Consistent with previous reports that the miR-9a-*sens* interaction in the epithelium modulates sensory neuron function (Li et al. 2006; Mullard 2006; Cassidy et al. 2013b; Gallicchio et al. 2020), our findings support a model where this regulatory axis is critical for shaping neuronal architecture. We conclude that the epithelial interaction between *miR-9a* and its target *sens* modifies the morphology of body wall sensory neurons, which in turn contributes to the regulation of complex behaviors like female receptivity.

### Spatial transcriptomics reveal an epithelial locus for *miR-9a* action

Through bioinformatic analysis, we identified several *miR-9a* target genes that contain *miR-9a* target sequences in their untranslated regions (UTRs) of mRNA (Kozomara et al. 2019). We conducted similar genetic interaction tests as with *sens* mutant and found that the removal of one copy of *bruno2* (*bru2/+*) completely rescued the *miR-9a* mutant receptivity phenotype (Fig. 5A). To determine the precise cellular locus of these regulatory interactions and to investigate why only a subset of predicted targets genetically modify the *miR-9a* phenotype, we analyzed publicly available *Drosophila* scRNA-seq data (Li et al. 2022). This allowed us to map the spatial expression patterns of *miR-9a* and its targets relative to *ppk*-expressing neurons.

**Figure 5.**
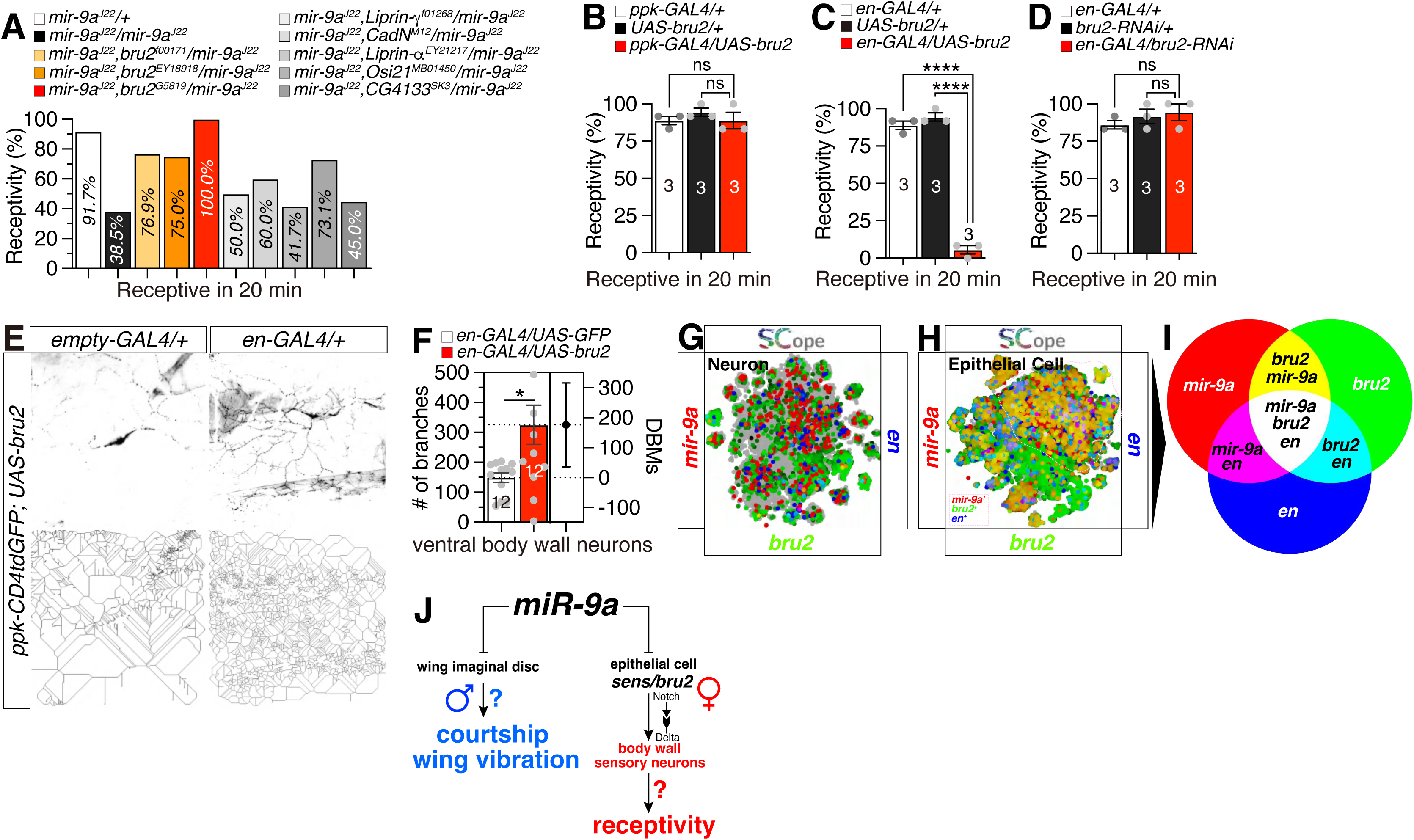
*miR-9a* guides sensory neuron development in the adult body wall and modulates virgin female receptivity through target gene regulation. (A-D) Receptivity of virgin females in 20 minutes, with the score for receptivity behavior as well as genotype of experimental animals given above the graph. (E) Overexpression of *bru2* in body wall sensory neurons via *en-GAL4* with *ppk-CD4tdGFP*. (F) Quantification of neurite morphology for body wall sensory neurons in branches. Genotype of experimental animals given above the graph. (G-H) Single-cell RNA sequencing (SCOPE scRNA-seq) datasets reveal tissues colored by expression of miR-9a (red), en (blue) with sens (green). (I) Schcematic diagram for color code presented in (G-H). (J) Conceptual model for the non-cell-autonomous role of *miR-9a* in modulating neurodevelopment and behavior. The schematic depicts a pathway where *miR-9a*, acting in epithelial cells, post-transcriptionally represses its targets sens and *bru2*. This repression in the epithelium provides a permissive environment for the correct dendritic patterning of neighboring *ppk^+^* neurons. Loss of *miR-9a* disrupts this intercellular signaling, causing neuronal overgrowth and impairing female receptivity, phenotypes that are rescued by reducing the dosage of *sens* or *bru2*.

Consistent with a non-cell-autonomous mechanism, we found that *miR-9a* is robustly expressed in epithelial cells but is largely absent from neuronal cell clusters marked by *ppk* expression (Fig. 3G-I). This spatial separation makes a direct, cell-intrinsic role for miR-9a in *ppk* neurons unlikely. We next examined the expression of the targets identified in our genetic screen (Fig. 5A). We found that *Liprin-α*, *Liprin-γ*, and *CadN* are expressed in both epithelial and neuronal tissues. However, a clear pattern emerged: in the epithelium, their expression domains showed significant overlap with miR-9a, whereas in neuronal clusters, they did not co-localize with *ppk* (Fig. S5A-I). In contrast, the predicted target *CG4133* was expressed predominantly in neurons, where its expression overlapped with miR-9a but not with *ppk* (Fig. S5M-O). Finally, another predicted target, *Osi21*, showed minimal expression in either epithelial or neuronal lineages in this dataset, suggesting it is not a primary target in this specific context (Fig. S5J-L).

Taken together, these transcriptomic data demonstrate a clear spatial segregation: *miR-9a* and its key genetic interactors, whose loss-of-function rescues the receptivity phenotype, are co-expressed in the epithelium (Supplementary Table 1). The affected *ppk*-positive neurons, however, constitute a separate cellular domain that lacks significant expression of these key targets. These findings provide strong transcriptomic support for a model in which *miR-9a* does not function within *ppk* neurons themselves but rather acts from the surrounding epithelium to control neuronal development and, consequently, female reproductive behavior.

### An epithelial *miR-9a-bru2* interaction non-cell-autonomously modulates sensory neuron function

Overexpression of *bru2* in sensory neurons did not alter the receptivity phenotype (Fig. 5B), indicating that *bru2* is not a direct target of *miR-9a* in sensory neurons. The *Drosophila* gene *bru2* is predicted to facilitate mRNA 3’-UTR binding activity and is involved in the negative regulation of translation (Öztürk-Çolak et al. 2024). Altered expression of *sens* can change sensory-organ specification (Li et al. 2006), and specific *ppk^+^* sensory neurons have been shown to control post-mating and egg-laying behaviors (Yapici et al. 2008; Häsemeyer et al. 2009b; Lee et al. 2016), whereas Bruno-family proteins influence oogenesis and flight-muscle maturation (Webster et al. 1997; Spletter et al. 2015), supporting the interpretation that egg-laying and wing-vibration phenotypes may arise indirectly. One of the identified targets, *sens*, is a well-established *miR-9a* target during sensory organ development (Li et al. 2006), and its expression in *ppk^+^* neurons is consistent with the strong morphological and behavioral effects we observed.

We next investigated other predicted targets of *miR-9a*, such as the RNA-binding protein *bru2*. While the function of *bru2* in *ppk^+^* neurons has not been characterized, potentially explaining the weak or absent phenotypes observed upon its dosage reduction, its role in female receptivity was evident in other tissues. Targeted overexpression of *bru2* in the epithelium using *en-GAL4* caused a significant reduction in female receptivity compared to controls. Conversely, RNAi-mediated knockdown of *bru2* in the same cells did not alter this behavior (Fig. 5C-D).

Epithelium-specific overexpression of *bru2* produced effects highly reminiscent of those obtained with sens. In ppk-positive body wall sensory neurons, this manipulation led to a significant increase in dendritic branching (Fig. 5E–F), recapitulating the morphological phenotype of the miR-9a mutant. Consistent with these phenotypic similarities, single-cell transcriptomic analysis revealed that miR-9a, bru2, and en are co-expressed within epithelial clusters but are absent from the neuronal groups expressing ppk (Fig. 5H–I, cf. Fig. 3G–I). This asymmetrical outcome is consistent with *bru2* functioning as a downstream target of *miR-9a* regulation. Collectively, these findings indicate that *miR-9a* guides sensory neuron specification and controls virgin female receptivity by fine-tuning the expression of multiple target genes, including sens and *bru2* (Fig. 5J).

To elucidate the precise cellular context of the *miR-9a-bru2* interaction, we analyzed their spatial expression patterns using the *Drosophila* single-cell transcriptomic atlas, Fly SCope (Li et al. 2022). The data revealed that *miR-9a* and its target, *bru2*, are robustly co-expressed in epithelial cells. In striking contrast, *bru2* transcript was found to be largely absent from the *ppk*-positive sensory neuron population (Fig. S5Q-S).

These distinct expression profiles preclude a direct, cell-autonomous role for *bru2* regulation by *miR-9a* within the *ppk*^+^ neurons themselves. Instead, this finding strongly supports a model of non-cell-autonomous regulation, where the molecular interaction occurs in one cell type (the epithelium) to influence the function of a neighboring cell type (the neuron). To further validate this regulatory relationship, we performed qRT-PCR analysis and found that *bru2* mRNA levels were significantly elevated in miR-9a homozygous mutants (Fig. S5P). We therefore propose that *miR-9a*-mediated repression of *bru2* within the epithelium indirectly modulates the physiological function of adjacent *ppk*^+^ sensory neurons.

Given its canonical role in mediating short-range cell-cell communication, particularly between epithelial and neural cells during development, we hypothesize that this non-cell-autonomous effect is propagated via the Notch-Delta signaling pathway (Artavanis-Tsakonas et al. 1999; Hori et al. 2013). This proposed mechanism, wherein the genetic interaction in epithelial cells influences neuronal function, is illustrated in our working model (Fig. 5J).

## DISCUSSION

Female *miR-9a* mutants display a pronounced phenotype characterized by increased rejection of courting males, delayed onset of sexual receptivity, and abnormal mating termination behavior. While the post-mating rejection behavior of mated females remains intact, they lay more eggs, suggesting specific effects on egg-laying (Fig. 1). Similar locomotor deficits are observed in both sexes, highlighting the generalized impact of *miR-9a* mutation on motor function. In male flies, *miR-9a* mutation leads to reduced courtship activity and the emergence of an abnormal bilateral wing vibration phenotype (Fig. 2). Further investigation of sensory neuron development reveals ectopic overgrowth of sensory neurons in both larval and adult stages, indicating a disruption in neuronal specification and growth (Fig. 3). Genetic interaction experiments with *sens* and *bru2*, genes involved in sensory bristle development and mRNA translation regulation, respectively, demonstrate that removing one copy of these genes in *miR-9a* mutant backgrounds rescues the receptivity defects and normalizes neuronal morphology (Fig. 4 and 5). As summarized in Fig. 5J, we propose a working model in which *miR-9a* influences sensory neuron morphology and virgin female receptivity, with *sens* and *bru2* acting as non-cell autonomous genetic modifiers. We note, however, that the roles of these genes in egg-laying and the function of *bru2* in body-wall sensory neurons remain to be tested, and thus the schematic should be viewed as a conceptual framework for future studies rather than a definitive pathway. These findings collectively suggest that *miR-9a* plays a crucial role in regulating the development of sensory neurons and female receptivity by modulating the expression of target genes such as *sens* and *bru2*. This study provides valuable insights into the molecular mechanisms underlying reproductive behaviors and neural development in *Drosophila*, and potentially in other organisms as well.

Although the role of miRNAs in neural development and plasticity is well-documented, their specific contributions to the modulation of sensory hypersensitivity following mating in female insects have not been extensively explored. Studies using genetic tools to manipulate miRNA expression in specific neurons or at specific times after mating could help to elucidate the precise mechanisms by which miRNAs contribute to these complex behavioral and physiological changes. Such studies are poised to elucidate the molecular underpinnings of post-mating sensory adaptations and their implications for reproductive success. Furthermore, the behavioral phenotypes observed in *miR-9a* mutants could be influenced not only by altered sensory processing but also by broader motor deficits. *miR-9a* has been implicated in neural development beyond reproductive circuits, and defects in locomotor performance have been reported in assays such as climbing. While our data suggest a strong correlation between sensory neuron overgrowth and changes in female receptivity, we cannot exclude the possibility that motor impairments may also contribute to the observed male courtship and female rejection behaviors. Future studies incorporating detailed analyses of motor circuits and electrophysiological characterization of sensory neuron excitability will be necessary to disentangle the relative contributions of sensory versus motor dysfunctions to these sexual behaviors.

While our data show overgrowth of *ppk^+^* body wall neurons in *miR-9a* mutants, the causal role of these neurons in female receptivity remains unresolved. Prior studies of *ppk^+^* neurons in the uterus and oviduct indicate that distinct subsets of *ppk^+^* neurons mediate post-mating responses (Häsemeyer et al. 2009b; Yang et al. 2009b; Rezával et al. 2012). Whether these subsets also overgrow in *miR-9a* mutants, whether body wall neurons are required for virgin receptivity, and whether their activity changes after copulation are key questions for future work. Addressing these points with cell-type–specific manipulations and functional imaging will be essential to clarify how *miR-9a* regulates *ppk^+^*neuronal subsets and reproductive behaviors.

In our investigation, we have elucidated a heretofore unreported target gene of *miR-9a*, designated *bru2*, which, upon deletion of a single allele, can rescue the female rejection phenotype induced by *miR-9a* mutation. The *Drosophila melanogaster bru2* gene is inferred to possess mRNA 3’-UTR binding activity and is proposed to participate in mRNA splice site recognition, the negative regulation of translation, and the regulation of alternative mRNA splicing, mediated through the spliceosome (Delaunay et al. 2004) . Despite these predictions, the precise molecular function of bru2 remains to be fully elucidated. The human orthologs of bru2, CELF1 (CUGBP Elav-like family member 1) and CELF2 (CUGBP Elav-like family member 2), have been implicated in developmental and epileptic encephalopathy 97 (Itai et al. 2021). Notably, a constellation of repressed expression of *let-7g*, *miR-9*, and *miR-135a*, alongside elevated expression of the RNA splicing factor CELF1, has been correlated with a murine heart model of myotonic dystrophy (MD) (Misra et al. 2020). These findings suggest that the newly characterized genetic interplay between *miR-9a* and *bru2* may serve as a prototypical model for investigating the role of human miR-9 in neuronal specification.

Our findings indicate that *miR-9a* mutations impact reproductive behaviors in both sexes, but with distinct outcomes. In females, *miR-9a* loss leads to reduced receptivity and increased rejection of courting males, correlating with sensory neuron overgrowth. In males, however, *miR-9a* mutants exhibit altered courtship performance, suggesting that *miR-9a* also contributes to the regulation of male-specific circuits underlying sexual display behaviors. These observations highlight that *miR-9a* exerts sex-specific effects on reproductive behavior, likely through its influence on neural development and circuit function in both male and female nervous systems. Considering these differences provides a more complete view of *miR-9a*’s role in behavioral modulation and raises important questions about how microRNAs contribute to sexually dimorphic neural and behavioral traits.

Our research revealed that mutation of *miR-9a* is associated with overgrowth of sensory neurons in adult *D. melanogaster* females, which correlates with increased rejection of courting males by virgins. While our results suggest a potential link between altered neuronal morphology and hypersensitivity-related behaviors, direct causal evidence of neuronal excitability changes remains to be established. The *miR-9a* mutant females exhibited a hypersensitivity phenotype attributable to this neuronal overgrowth. The study of hypersensitivity in *D. melanogaster* is a field of inquiry that holds profound implications for comprehending the foundational mechanisms underpinning sensory perception and neural plasticity. Although we did not perform *ppk*-restricted *miR-9a* rescue or *ppk*-specific *miR-9a* knockdown in this study, multiple lines of genetic evidence support a substantive role for *ppk^+^* body-wall sensory neurons: dendritic overgrowth maps to *ppk^+^* cells, and reducing *sens* or *bru2* dosage rescues both morphology and receptivity. We therefore view targeted re-expression and tissue-specific loss-of-function of *miR-9a* as important next steps to test cell autonomy and temporal requirements.

Hypersensitivity in sensory neurons can arise from changes in neural development or plasticity, leading to exaggerated responses to normal stimuli (Petersen-Felix and Curatolo 2002; Latremoliere and Woolf 2009a; Gangadharan and Kuner 2013). This can involve altered gene expression in sensory neurons, changes in the morphology and function of these neurons, and modifications in the way they interact with other cells in the nervous system (Woolf and Salter 2000). In some cases, hypersensitivity can become chronic due to central sensitization, a process where the central nervous system becomes hyper-excitable and develops an increased sensitivity to incoming sensory signals. This can result in long-lasting pain even after the initial injury or inflammation has resolved (Salter 2010).

Our study reveals the critical role of *miR-9a* in regulating reproductive behaviors and neural development in *Drosophila melanogaster*. Sensory hypersensitivity refers to an exaggerated response to normal stimuli, often manifesting as pain or discomfort in contexts where such sensations would not typically occur (Latremoliere and Woolf 2009b; Isaacs and Riordan 2020). It is frequently linked to neurological disorders, injury, or inflammation (Ren and Dubner 2008; Pinho-Ribeiro et al. 2017). Understanding the mechanisms that drive hypersensitivity in sensory neurons is therefore critical for insights into both normal sensory processing and pathological conditions.

In *D. melanogaster*, post-mating modifications in female behavior encompass enhanced sensory reactivity, a phenomenon considered adaptive as it facilitates the evasion of predators and the selection of appropriate oviposition sites (Chapman et al. 2003; Ram and Wolfner 2007; Yapici et al. 2008; Gligorov et al. 2013; Corbel et al. 2022). This augmented sensitivity is correlated with the overgrowth of adult body wall sensory neurons, and alterations in *miR-9a* expression via genetic manipulations can modulate these responses. The analysis of hypersensitivity in this fly model provides critical insights into the molecular and neural substrates that regulate sensory processing and the evolution of complex behavioral outputs. Our study establishes a correlation between sensory neuron overgrowth and altered virgin female receptivity, but it does not prove causation. It remains possible that an upstream regulatory event independently affects both neuronal morphology and behavior. Future experiments combining electrophysiology with cell-specific manipulations will be essential to determine whether structural changes in body wall neurons directly underlie behavioral phenotypes.

Notably, female *ppk*-positive body wall neurons have been identified as key neurons underlying the heat hypersensitivity phenotype (Gu et al. 2022), suggest that adult body wall neurons contribute to adaptive hypersensitivity, consistent with conserved roles of sensory neurons in mediating heightened sensitivity across species. Our discovery establishes a novel *Drosophila* genetic model for the investigation of *miR-9* family-mediated pathogenesis across various neuronal contexts.

Our research identifies a critical role for *miR-9a* in regulating female receptivity, a complex behavior in *Drosophila* that is contingent on the proper morphological development of body wall sensory neurons. The central finding of our work is that this regulation occurs through a non-cell-autonomous mechanism. We propose a model where *miR-9a* acts not within the neuron itself, but within the surrounding epithelial cells, to orchestrate a microenvironment that is permissive for correct neuronal dendritic growth. This finding shifts the understanding of *miR-9a*’s role from a direct, intracellular regulator of neuronal architecture to a master regulator of the intercellular signaling that underpins neurodevelopment.

The strongest evidence supporting this model comes from our genetic epistasis and phenocopy experiments. While the loss of *miR-9a* leads to dendritic overgrowth in *ppk* neurons and a corresponding decrease in female receptivity, we demonstrate that this phenotype is not due to a primary defect within the neuron. Instead, the phenotype can be fully recapitulated by overexpressing *miR-9a*’s downstream targets—the transcription factor *sens* or the RNA-binding protein *bru2*—specifically within the epithelium. This crucial result, where manipulating a gene in one cell type (epithelium) produces a phenotype in an adjacent cell type (neuron), is the definitive signature of non-cell-autonomous action. It strongly implies that the primary function of *miR-9a* in this context is to repress sens and bru2 within the epithelium. In the absence of miR-9a, these targets become derepressed, altering the signaling properties of epithelial cells and leading them to send aberrant growth cues to the neighboring neurons.

Our model posits that the epithelial derepression of *sens* and *bru2* is the key molecular event that disrupts epithelial-neuronal communication. *Sens* is a well-characterized transcription factor essential for peripheral nervous system development, particularly in specifying the fate of sensory organ precursors (Nolo et al. 2000; Li et al. 2006; Mullard 2006; Cassidy et al. 2013b; Gallicchio et al. 2020). Its misexpression in the broader epithelial tissue could fundamentally alter the identity of these cells, causing them to adopt signaling characteristics that pathologically promote neuronal growth. *Bru2*, as an RNA-binding protein involved in translational repression, likely contributes by altering the proteome of epithelial cells (Chekulaeva et al. 2006). The derepression of *bru2* could lead to the inappropriate translation of a suite of mRNAs, including those encoding secreted ligands, cell-surface receptors, or components of the extracellular matrix that collectively influence neuronal guidance and growth.

The coordinated misregulation of a transcription factor (*sens*) and an RNA-binding protein (*bru2*) highlights the sophisticated regulatory logic of miRNAs. Rather than targeting a single linear pathway, *miR-9a* acts as a nodal point to simultaneously control gene expression at both the transcriptional and post-transcriptional levels. This allows for a robust and multi-faceted regulation of the epithelial cell’s signaling output. This work, therefore, adds to a growing body of evidence that the epidermis is not merely a passive scaffold for the nervous system but an active and indispensable signaling center that sculpts neuronal morphology, a concept well-established in the context of dendritic tiling and boundary formation (Li et al. 2006; Mullard 2006; Jones 2008; Bartel 2009b; Biryukova et al. 2009; Kadener et al. 2009; Smalheiser and Lugli 2009b; Cohen et al. 2011b; Yuva-Aydemir et al. 2011b; Cassidy et al. 2013b; Aksoy-Aksel et al. 2014b; Søvik et al. 2015; Wang et al. 2016b; Alberti et al. 2018; Ridler 2018; Xue and Zhang 2018; Mohammadi et al. 2022b).

This non-cell-autonomous model provides a robust framework for understanding the link between miR-9a and female receptivity, but it also opens several exciting avenues for future investigation. The most immediate question is the identity of the downstream signaling molecule(s) acting between the epithelium and the neuron. The aberrant epithelial signals initiated by *sens* and *bru2* overexpression could be secreted ligands, cell-adhesion molecules, or changes in the extracellular matrix such as Notch-delta interaction (Li et al. 2006). An unbiased transcriptomic or proteomic screen comparing wild-type and *miR-9a* mutant epithelial tissue could identify candidate signaling factors. Furthermore, our study primarily focuses on the developmental role of this pathway. It remains an open and intriguing question whether *miR-9a* continues to function in the adult epithelium to maintain the sensory circuit’s integrity and function. A conditional, adult-stage-specific knockdown of *miR-9a* or its targets in the epithelium would be a powerful experiment to dissect any ongoing role of this pathway in neuronal maintenance and behavioral plasticity. Answering these questions will not only illuminate the specific mechanism of *miR-9a* function but also provide deeper insights into the fundamental principles governing tissue-level communication in the development and maintenance of a functional nervous system.

## METHODS

### Fly Stocks and Husbandry

*Drosophila melanogaster* were raised on cornmeal-yeast medium at similar densities to yield adults with similar body sizes. Flies were kept in 12 h light: 12 h dark cycles (LD) at 25°C (ZT 0 is the beginning of the light phase, ZT12 beginning of the dark phase) except for some experimental manipulation. Wild-type flies were *Canton-S* (*CS*) and *Cantonized w1118*. Following lines used in this study, *Canton-S* (#64349), *ppk-CD4-tdGFP* (# 35842), *sens^E58^* (#5312), *bru2^f00171^* (#18300), *bru2^EY18918^* (#22296), *bru2^G5819^* (#27190), *Liprin-γ ^f01268^* (#18421), *CadN^M1^*^2^ (#229), *Liprin-α^EY21217^* (#22459), *Osi21^MB0145^ ^0^*(#23186), *ppk-GAL4* (#32078), *en-GAL4* (#1973) were obtained from the Bloomington *Drosophila* Stock Center at Indiana University. *Osi21^MB01450^* (#M2L-3116) was obtained from National Institute of Genetics Fly Stocks. We thank Dr. Fen-Biao Gao (Gladstone Institute of Neurological Disease and Department of Neurology, University of California at San Francisco) and Dr. Kweon Yu (KRIBB) for sharing *miR-9a^J22^*, *miR-9a^E39^*, and *UAS-miR-9a* lines. We thank Dr. Paul M. Macdonal (Department of Molecular Biosciences, Institute for Cellular and Molecular Biology, The University of Texas at Austin, Austin, Texas, United States of America) for sharing *UAS-bru2* line. We thank Dr. Hugo Bellen (Baylor College of Medicine) for sharing *UAS-sens* line.

### Receptivity Assay and Egg Laying Assay

For receptivity assays, females and males were housed in small groups of 3–4 flies. In order to assess female receptivity, one female was put into a small (1 cm × 1 cm) chamber together with two naive *Canton S* males. The female was scored as receptive if it mated within 20 min. In the remating assays, a female was mated in a receptivity assay and then again examined in another receptivity assay 24 hours later. Assays were usually performed within the first three hours of the experiment day. Rejection (%) was calculated as the total number of rejection behaviors observed (e.g., abdominal turning, kicking) during the observation period divided by the number of tested females ×100. Because a single female could display multiple rejection behaviors, the rejection percentage could exceed 100%. All experimental procedures were conducted using 36-well plates, and each assay was replicated the specified number of times to ensure statistical robustness. The calculated percentages of receptivity and rejection were then subjected to statistical analysis via Student’s t-test for significance.

For egg laying assays, 10 mated females of the appropriate genotype were aged in vials for 4–5 days. Then three or five (as indicated in the figure legends) females were transferred to a vial with grape media and allowed to lay eggs for 24 hours at 25°C. The number of eggs was divided by the number of flies in the vial to give a measure of egg laying. For assays of egg laying by mated females, females of the respective genotype were mated with *Canton S* males for 2–3 hours on the day before the experiment, with 10 females and 20 males per vial. All data are given as average ± SEM, significance levels were calculated with the Student’s t test.

### Climbing Assay

For climbing assay, we modified the conventional RING assay (Gargano et al. 2005) and reported in our previous reports (Miao et al. 2024; Zhang et al. 2024). In brief, 40-50 aged flies were placed in an empty vial and were tapped to the bottom of the tube. We used 5 days old adults. After tapping of flies, we recorded 10 seconds of video clip. This experiment was repeated twice for each group at 5-minute intervals. For analysis, the performance values from the two trials were averaged for each group to obtain a single data point, which was used in statistical comparisons. With recorded video files, we captured the position of flies 10 seconds after tapping the vial. This captured image file was then loaded in ImageJ to perform particle analysis. For quantifying the location of flies inside a vial, we used the “analyze particles” function of ImageJ (Grishagin 2015). The position of pixels was normalized by height of vial then only the particles above the midline (4 cm) of vial were counted.

### Adult Flight Assay

Flies were subjected to a ’flight assay’ to evaluate their escape response from water. 50 flies were gently introduced into a water-filled jar. The jar was then tapped to stimulate the flies, and the number of flies that escaped from the water was counted. The escape ratio was calculated to determine the effectiveness of the flies’ flight response. For visual reference, please check the video files labeled 5-6, which represent the phenotypes of control and mutant flies during the assay. Our experiment was conducted at 25°C under normal light conditions.

### Courtship Assay

The courtship assay was conducted according to established protocols as previously reported (Lee et al. 2023), under standard light conditions in circular courtship arenas with a diameter of 11 mm, between noon and 4 p.m. Courtship latency was defined as the interval from the introduction of the female to the initiation of the first overt male courtship behavior, such as orientation and wing extensions. Following the onset of courtship, the courtship index was determined as the proportion of time the male engaged in courtship-related activities over a 10-minute period or until copulation occurred.

### Adult Body Wall Neuron Live Imaging

Adult body wall was dissected and fixed in 4% paraformaldehyde in PBS for 30–60 min at room temperature, followed by blocking for 30 min in PBS containing 0.3% Triton X-100 and 5% normal goat serum. Tissues were stained with rat anti-mCD8 (1:200; Caltag) and/or mouse anti-armadillo (DSHB; N2 7A1, 1:15). Primary antibodies were detected with Cy2-conjugated goat anti-rat or Cy5-conjugated goat anti-mouse secondary antibodies (Jackson Research). LTM fibers were labeled with phalloidin-TRITC (1:200; Sigma) or Alexa647-phalloidin (1:200; Invitrogen). Images were taken on a Leica TCS SP5 confocal microscope (Leica). As an alternative to antibody staining, we imaged GFP and mCherry fluorescence in living animals by mounting them in silicon oil (Shin-Etsu). Maximum projections of z-stacks were used in all cases. Images were adjusted for brightness and contrast with Adobe Photoshop (Adobe Systems, San Jose, CA) (Yasunaga et al. 2015). For visualization of dendrites, we labeled neurons with ppk-GAL4; UAS-mCD8GFP, ppk-CD4-tdGFP, or ppk-CD4-tdTom and imaged GFP and RFP fluorescence in living animals by mounting them in silicon oil (Shin-Etsu). Maximum projections of Z-stacks were used in all cases. The dendrite length was measured by using the ImageJ. For the quantitative analyses, we focused on the dendrites in segments A4, A5 and A6, since these neurons exhibit similar and consistent dendrite branch lengths and branch points. For quantification of the total branch length and the branch points, we used skeleton analysis as described above.

### Quantification of Neurite Branching

To quantify neurite branching, we utilized the ’skeleton’ function in ImageJ, following a detailed step-by-step process to ensure accurate measurement of branch numbers and lengths. All specimens were imaged under identical conditions. First, images of neurons were imported into ImageJ by launching the software and dragging the image into the workspace. The images were then converted to an 8-bit format by selecting the image, navigating to the “Image” menu, choosing “Type,” and selecting “8-bit.” Next, the images underwent threshold adjustment to separate the neuron branches from the background. This involved selecting “Image” from the menu bar, choosing “Adjust,” and then “Threshold.” The “Dark Background” option was checked, the color was changed to red, and the threshold slider was adjusted until all branches were highlighted in red. After threshold adjustment and branch connection, the images were converted to a binary format. This was done by selecting “Process” from the menu bar, choosing “Binary,” and then “Make Binary,” ensuring that “Threshold pixel to foreground color” and “Remaining pixel to background color” options were checked with a “Black foreground, white background” setting. The binary images were then skeletonized to produce a simplified representation of the neurite branches. This involved selecting “Process” from the menu bar, choosing “Binary,” and then “Skeletonize.” The resulting skeleton images were analyzed using the Analyze Skeleton plugin. This process included selecting “Analyze” from the menu bar, choosing “Skeleton,” and then “Analyze Skeleton.” The settings for the analysis included keeping the “Prune cycle method” set to “None,” unchecking the “Prune ends” and “Exclude ROI from pruning” options, and checking the options to show the longest shortest path, branch labels, junctions, and endpoints. All specimens were imaged under identical conditions. In our skeleton analysis, a branch was defined as a continuous dendritic segment between either a junction and an endpoint or between two junctions. A junction was defined as a branch point where one dendrite splits into two or more daughter dendrites. These definitions were applied consistently across all genotypes. We also counted the total number of sensory neurons in the imaged abdominal segments (A4–A6) to assess whether neuron number was altered.

### Predicting of *miR-9a* Targets in *Drosophila* Genome using bioinformatical anlaysis

We used miRBase provided targetscan analysis to identify *miR-9a* potential targets (Kozomara et al. 2019). The result can be checked with this stable link. https://www.targetscan.org/cgi-bin/targetscan/fly_12/targetscan.cgi?mirg=dme-*miR-9a*

### Video Recording of Larval Locomotor Behaviors

Video recordings of gross path morphology were made with a digital video (DV) camera (Canon GL1) in an environment room maintained at 25°C and 70% humidity. DV movies were captured with IMOVIE 2.0 on a 500-MHz Apple iMac and digitized with QUICKTIME 4.0 at 29.97 frames per second (fps). Low-magnification videos were recorded for 2 min or until the larva left the 50-cm^2^ field of the camera. High-magnification videos (15-cm^2^ field) were recorded with an Olympus OLY-200 camera mounted on an Olympus SZX9 microscope connected to a VCR (Samsung VR5599). Peristalsis was recorded until 10 peristaltic waves during linear locomotion were completed. Mutants frequently had discontinuous bouts of peristalsis, and repositioning of the plate was necessary, but this did not affect the motion analysis. VCR recordings were captured with adobe premier 5.0 on a Macintosh G4 at 15 fps.

### Quantitative real time PCR (qRT-PCR)

The expression levels of *bru2* gene in wild-type and *miR-9a* mutant female flies in different conditions were analyzed by quantitative real time PCR (qRT-PCR) with SYBR Green qPCR MasterMix kit (Selleckchem). Primers for amplifying the genes in qRT-PCR were designed following previous reports (Boutros et al. 2004; Hu et al. 2013), F: 5’–AAATTCGCCGACACGCAAAA-3’; R: 5’–CCATCGACGGATTGGTACGT–3’. qPCR reactions were performed in triplicate, and the specificity of each reaction was evaluated by dissociation curve analysis. Each experiment was replicated three times. PCR results were recorded as threshold cycle numbers (Ct). The fold change in the target gene expression, normalized to the expression of internal control gene (GAPDH) and relative to the expression at time point 0, was calculated using the 2 −ΔΔCT method as previously described (Livak and Schmittgen 2001). The results are presented as the mean ± SD of three independent experiments.

### Statistical Analysis

Statistical analysis of receptivity assays is similar with mating duration assay was described previously (Lee et al. 2023). More than 10 females for 1 group were used for receptivity assay. Statistical comparisons were made between groups that were control group and experimental group within each experiment. As receptivity assays of females showed normal distribution (Kolmogorov-Smirnov tests, p > 0.05), we used two-sided Student’s t tests. The mean ± standard error (s.e.m) (*\*\*\*\* = p < 0.0001, *** = p < 0.001, ** = p < 0.01, * = p < 0.05*). All analysis was done in GraphPad (Prism). Individual tests and significance are detailed in figure legends. Besides traditional t-test for statistical analysis, we added estimation statistics for all receptivity assays and two group comparing graphs. In short, ‘estimation statistics’ is a simple framework that—while avoiding the pitfalls of significance testing—uses familiar statistical concepts: means, mean differences, and error bars. More importantly, it focuses on the effect size of one’s experiment/intervention, as opposed to significance testing (Claridge-Chang and Assam 2016). For DBMs (Difference Between Means) plots, the central dot represents the mean difference, and the error bars represent the 95% confidence interval, computed by bootstrap resampling in the estimation statistics framework. In comparison to typical NHST plots, estimation graphics have the following five significant advantages such as (1) avoid false dichotomy, (2) display all observed values (3) visualize estimate precision (4) show mean difference distribution. And most importantly (5) by focusing attention on an effect size, the difference diagram encourages quantitative reasoning about the system under study (Ho et al. 2019). Thus, we conducted a reanalysis of all of our two group data sets using both standard t tests and estimate statistics. In 2019, the Society for Neuroscience journal eNeuro instituted a policy recommending the use of estimation graphics as the preferred method for data presentation (Bernard 2021).

## DATA AVAILABILITY STATEMENT

The authors affirm that all data necessary for confirming the conclusions of the article are present within the article, figures, and tables.

## ACKNOWLEDGEMENTS

We are very appreciative to the colleagues who supplied us with several fly strains. We thank Dr. Fen-Biao Gao (Gladstone Institute of Neurological Disease and Department of Neurology, University of California at San Francisco) and Dr. Kweon Yu (KRIBB) for sharing *miR-9a^J22^*, *miR-9a^E39^*, and *UAS-miR-9a* lines. We thank Dr. Paul M. Macdonal (Department of Molecular Biosciences, Institute for Cellular and Molecular Biology, The University of Texas at Austin, Austin, Texas, United States of America) for sharing *UAS-bru2* line. We thank Dr. Hugo Bellen (Baylor College of Medicine) for sharing *UAS-sens* line. We extend our gratitude to Drs. Yuh Nung and Lily Jan (UCSF) for their invaluable support and engagement in discussions pertaining to this project. This research was supported by a Startup funds from HIT Center for Life Science to WJK.

## DECLARATION OF GENERATIVE AI AND AI-ASSISTED TECHNOLOGIES IN THE WRITING PROCESS

During the creation of this work, the author(s) utilized DeepSeek AI (https://chat.deepseek.com/) to rephrase English sentences and verify English grammar, as none of the authors of this paper are native English speakers. After using this tool/service, the author(s) reviewed and edited the content as needed and take(s) full responsibility for the content of the publication.

Figure S1. *miR-9a* mutations lead to a specific increase in rejection behavior among virgin females.

(A-B) Receptivity of virgin females in 20 minutes, with the score for receptivity behavior as well as genotype of experimental animals given above the graph. (C) The temporal pattern of virgin female receptivity to males courtship across the duration of the observation period. (D) Rejection of mated females, with the score for rejection behavior as well as genotype of experimental animals given above the graph. (E) Egg laying. For each assay, five females of the indicated group were allowed to lay eggs in a vial with grape media at 25℃. Eggs were counted after 24 hours and the number of eggs in each vial was divided by five. n=15 assays for all genotypes, error bars indicate SEM, ∗p < 0.05, ∗∗∗p < 0.0005, Student’s t test.

Figure S2. *miR-9a* mutants exhibit locomotor defects among larvae and adult.

(A) Climbing assays. (B) Flight assays of females. For each assay, fifty flies were gently introduced into a water-filled jar. The jar was then tapped to stimulate the flies, and the number of flies that escaped from the water was counted. The escape ratio was calculated to determine the effectiveness of the flies’ flight response. (C) 3^rd^ instar larvae of *miR-9a^J22^/+* fly. (D) 3^rd^ instar larvae of *miR-9a^J22^/miR-9a^J22^* fly. (E) Speed of 3^rd^ instar larvae crawling. (F) Turn number of 3^rd^ instar larvae crawling.

Figure S3. miR-9a guides sensory neuron development in the adult and larvae dorsal body wall.

(A) Location of *Drosophila* larvae ventral body wall neurons. (B) Ventral neurons expressing *ppk-GAL4* together with *UAS-mCD4GFP* in *miR-9a^J22^/ +* and *miR-9a^J22^/miR-9a^J22^* in larvae. Scalebar represent 50 μm. (C) Location of *Drosophila* adult dorsal neurons. (D) Dorsal A5 neurons expressing *ppk-GAL4* together with *UAS-mCD4GFP* in *miR-9a^J22^/ +* and *miR-9a^J22^/miR-9a^J22^* in adult female. The bottom pictures are skeletonized from the top pictures. Scalebar represent 100 μm. (E-G) Quantification of neurite morphology for dorsal body wall neurons in branches (C), junctions (D) and branch length (E). Genotype of experimental animals given above the graph.

Figure S4. *sens^E58^/+* rescued the receptivity defects in a *miR-9a^J22^* homozygous mutant background.

(A) Receptivity of virgin females in 20 minutes, with the score for receptivity behavior as well as genotype of experimental animals given above the graph. (B) *ppk+* sensory neurons in the abdominal body wall (A4–A6) expressing *ppk-CD4tdGFP* in *wCS* and *miR-9a^J22^/miR-9a^J22^* in adult female.

Figure S5. **Epithelial localization of miR-9a targets supports non-cell-autonomous regulation.** (A-O) SCOPE scRNA-seq datasets and schcematic diagram reveal tissues colored by expression of *miR-9a* (red), *ppk* (green) with (blue) (A-C) *Liprin-γ*, (D-F) *CadN*, (G-I) *Liprin-α*, (J-L) *Osi21*, and (M-O) *CG4133*. (P) The results of qRT-PCR for *bru2* gene expression. Genotype of experimental animals given above the graph. The y-axis depicts the relative expression level of SIFaR, normalized to the expression of the GAPDH gene. “Rltv. Gene Exp” denotes relative gene expression. (Q-R) SCOPE scRNA-seq datasets reveal tissues colored by expression of *miR-9a* (red), *ppk* (blue) with *bru2* (green). (S) Schcematic diagram for color code presented in (Q-R).

## Supplementary Movie Descriptions

Movie 1-2. Receptivity in wild-type females.

A wCS virgin female is courted by wCS males. The female accepts the male and copulation proceeds normally. This represents typical receptivity in controls (related to Fig. 1A–B).

Movie 3-4. Rejection behavior in miR-9a mutant females.

A virgin *miR-9a^J22^/miR-9a^J22^* female is courted by wCS males. The female repeatedly rejects the courting males by turning and kicking. This illustrates the elevated rejection phenotype (related to Fig. 1D).

Movie 5. Flight assay in control flies.

Fifty Canton-S adults are tapped into a water-filled jar. Most rapidly escape by flying, reflecting normal flight ability (related to Fig. 1I).

Movie 6. Flight assay in miR-9a mutant flies.

Fifty *miR-9a^J22^/miR-9a^J22^* adults are tapped into a water-filled jar. Many fail to escape, reflecting impaired flight ability (related to Fig. 1I).

Movie 7. Courtship display in miR-9a mutant males.

A *miR-9a^J22^/miR-9a^J22^* male courts a Canton-S virgin female using abnormal bilateral wing vibration instead of the typical unilateral display. This phenotype is quantified in Fig. 2B, D.

Movie 8. Courtship display in control males.

A Canton-S male courts a virgin female using normal unilateral wing vibration. This serves as a control for Movie 7 (related to Fig. 2C, D).

Movie 9. Locomotion in control larvae.

A Canton-S third instar larva crawls forward smoothly with normal peristaltic waves. This represents typical larval locomotor behavior (related to Fig. S2C–E).

Movie 10. Locomotion in miR-9a mutant larvae.

A *miR-9a^J22^/miR-9a^J22^* third instar larva exhibits impaired locomotion with increased turning and disrupted peristalsis. This illustrates the larval locomotor phenotype (related to Fig. S2E–F).

